# PromptBio: A Multi-Agent AI Platform for Bioinformatics Data Analysis

**DOI:** 10.1101/2025.07.05.663295

**Authors:** Minzhe Zhang, Wenhao Gu, Bowei Han, Vincent Guo, Chintan Addoni, Jiayu Chen, Youjia Ma, Yang Leng, Kai Li, Xiaoxi Lin, Shi Shi, Junbin Zheng, Yilin Zheng, Weiying Wang, Linlin Wu, Linglang Yu, Juan Wang, KC Shashidhar, Xiao Yang

## Abstract

PromptBio is a modular AI platform for scalable, reproducible, and user-adaptable bioinformatics analysis, powered by generative AI and natural language interaction. It supports three complementary modes of analysis designed to meet diverse research needs. PromptGenie is a multi-agent system that enables stepwise, human-in-the-loop workflows using prevalidated domain-standard tools. Within PromptGenie, specialized agents—including DataAgent, OmicsAgent, AnalysisAgent, and QAgent—collaborate to manage tasks such as data ingestion, pipeline execution, statistical analysis, and interactive summarization. DiscoverFlow provides integrated, automated workflows for large-scale multi-omics analysis, offering end-to-end execution and streamlined orchestration. ToolsGenie complements these modes by dynamically generating executable bioinformatics code for custom, user-defined analyses, enabling flexibility beyond standardized workflows. PromptGenie and DiscoverFlow leverage a suite of domain-specific tools, including Omics Tools for standardized omics pipelines, Analysis Tools for downstream statistical interpretation, and MLGenie for machine learning and multi-omics modeling. We present the design, capabilities, and validation of these components, highlight their integration into automated and customizable workflows, and discuss extensibility, monitoring, and compliance. PromptBio aims to democratize high-throughput bioinformatics through a large language model–powered, natural language understanding, workflow generation and agent orchestration.

## 1 Introduction

Bioinformatics data analysis requires processing high-dimensional, heterogeneous datasets, such as genomics, transcriptomics, proteomics, and biomedical literature. Traditional pipelines often lack flexibility and scalability, prompting the development of AI-driven solutions. The PromptBio platform addresses this need through a modular, multi-agent system powered by generative AI, enabling automated, interpretable, and AI-human collaborative bioinformatics analysis. This paper presents the design and implementation of PromptBio, which coordinates specialized AI agents across data management, analysis, prediction, summarization, and interpretation tasks.

The advent of large language models (LLMs) is transforming bioinformatics by enabling natural language–driven workflows that reduce technical barriers and simplify complex analyses [1]. Incumbent platforms like DNAnexus [2] and Seqera [3] have advanced bioinformatics infrastructure but remain dependent on manual configuration and domain-specific expertise. In contrast, emerging AI-first platforms—such as Biomni [4], FutureHouse [5], Kepler AI [6], and Potato AI [7]— offer flexible, intelligent systems that broaden access to data-driven discovery. PromptBio (https://promptbio.ai) contributes to this evolution by offering an industrial-grade, agent-based platform that delivers value through: (1) making data analysis intuitive and engaging, (2) empowering users across technical skill levels, and (3) providing comprehensive, cost-effective solutions (https://promptbio.ai/use-cases.html).

Designing next-generation bioinformatics platforms demands adherence to several key principles. First, conversational interfaces driven by LLMs promote accessibility and ease of use [8]. Second, modular, multi-agent design enables scalable, parallelized workflows for tasks like multi-omics integration [1]. Third, robust evaluation protocols ensure scientific validity and reproducibility [9]. Fourth, real-time monitoring supports fault detection and system optimization [10]. Fifth, cloud-native infrastructure supports scalability and cost-efficiency [11], while strong security and regulatory compliance (e.g., HIPAA) ensure data protection [12]. Finally, extensibility allows the platform to adapt to emerging omics technologies and evolving analysis needs.

PromptBio distinguishes between three complementary modes of analysis:

(1) PromptGenie – a multi-agent system for managing and processing omics data, machine learning tasks, and tabular data operations. It leverages domain-standard, prevalidated tools with embedded best practices and emphasizes human-in-the-loop design to ensure precision and reproducibility.
(2) DiscoverFlow – an orchestration utility that streamlines complex workflows and supports end-to-end execution. DiscoverFlow enables automated, large-scale multi-omics analyses, facilitating the integration and management of diverse analytical components within a unified framework.
(3) ToolsGenie – a generative engine that empowers users to design and execute custom analyses tailored to specific research questions. ToolsGenie provides the flexibility needed for exploratory and innovative workflows, complementing standardized approaches with user-driven adaptability.

The platform’s intuitive, conversation-driven interface allows users—both expert and novice—to execute sophisticated bioinformatics tasks such as data preprocessing, clustering, differential analysis, and biomarker discovery without writing code. PromptBio’s multi-agent structure functions as a virtual interdisciplinary team, integrating expertise across data science, biology, and clinical research into a cohesive analytical unit. This design reduces communication overhead, accelerates analysis, and enables individual users to achieve team-level productivity.

In this paper, we describe the platform’s design, core tools, workflow orchestration, and operational safeguards. Section 2 introduces the multi-agent design. Section 3 presents the suite of built-in tools, including Omics Tools, Analyses Tools, MLGenie, and MarkerGenie. Section 4 describes DiscoverFlow, a framework for orchestrating automated multi-omics workflows. Section 5 discusses ToolsGenie, which supports on-demand, user-defined analyses via generative AI. Section 6 details platform monitoring, compliance, and security, and Section 7 outlines future directions toward fully agentic AI systems.

## 2 Multi-Agent Design

PromptBio’s multi-agent system (Figure 1) consists of a supervisor agent, PromptGenie, which coordinates a list of specialized agents—including the **OpsAgent, DataAgent, OmicsAgent, AnalysisAgent, and QAgent**—that collaboratively manage the full lifecycle of omics and machine learning tasks. These agents rely on prevalidated tools and domain-standard practices to ensure analysis accuracy, reproducibility, and interpretability.

**Figure 1:**
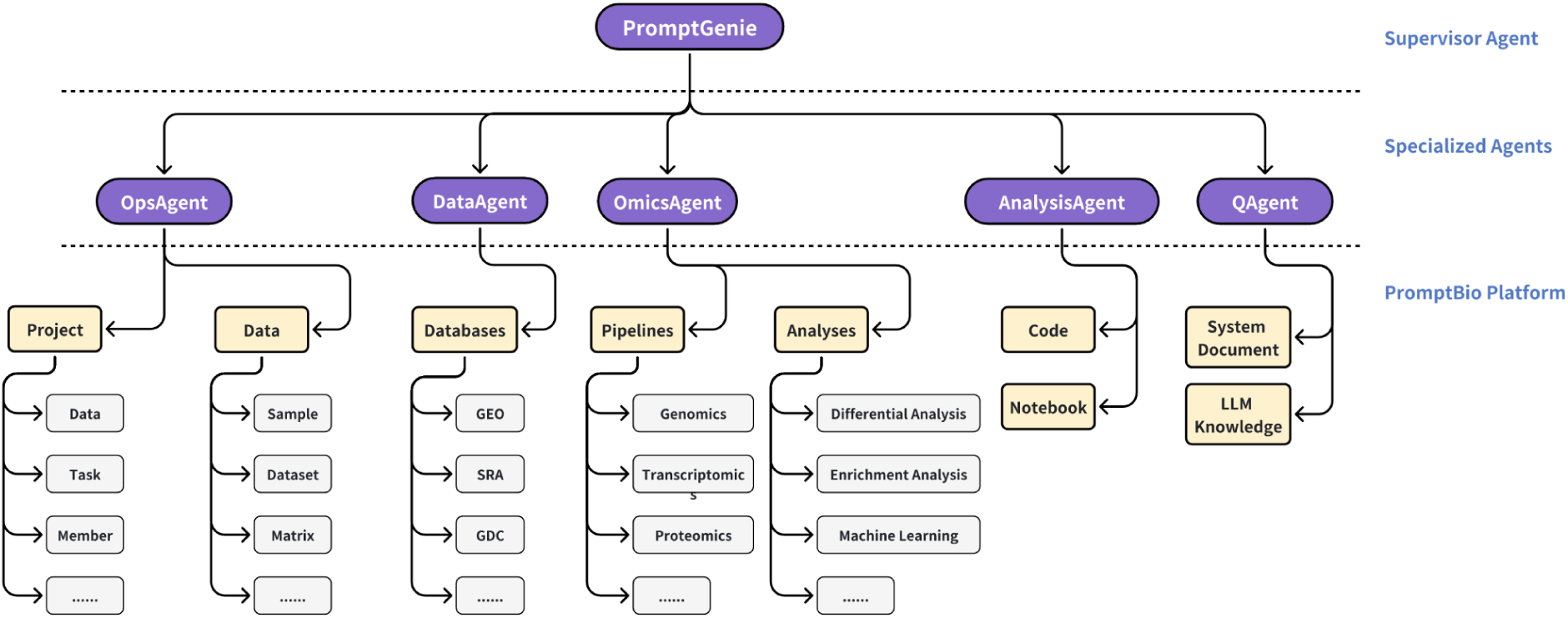
Design of the PromptBio multi-agent system, depicting interactions between PromptGenie and specialized agents, including OpsAgent, DataAgent, OmicsAgent, AnalysisAgent, and QAgent, and integrated tools.

Built on a modular and extensible design, PromptGenie enables the integration of new tools and supports task decomposition for scalable, parallel execution. Importantly, it incorporates a human-in-the-loop design, enabling domain experts to review intermediate steps, guide decision points, and validate outputs.

### 2.1 PromptGenie

PromptGenie, the supervisor agent, defines standard communication protocols, unified data schemas, and a shared runtime environment to ensure seamless operation across specialized agents.

At its core is an Agent Registry—a centralized directory that maps analytical tasks to appropriate agents (e.g., DataAgent, OmicsAgent) based on their declared capabilities. This enables targeted and efficient task routing.

User queries are submitted in natural language and interpreted by integrated LLMs, which translate them into executable steps. These steps are then routed to agents for immediate execution in a human-in-the-loop process that enables guided, interactive refinement. For example, commands such as “*Apply RNA-seq analysi*s” and “*Find differentially expressed genes*” are processed step by step, ensuring transparency, control, and reproducibility.

By integrating agentic execution with natural language interfaces, PromptGenie streamlines bioinformatics workflows in a way that is accessible to novices and powerful enough for experts—ensuring scalability, reproducibility, and ease of use within a structured yet adaptable framework.

### 2.2 OpsAgent

OpsAgent serves as the controller for data management, enforcing strict structural and relational integrity across all core entities—samples, subjects, datasets. It exposes a transactional interface for CRUD (create, read, update, delete) operations with atomicity guarantees and versioned audit trails, ensuring all modifications are consistent and traceable. This coordination abstracts away low-level data logistics, reducing operational overhead and minimizing the risk of duplication or corruption in collaborative environments—providing a robust foundation for scalable, compliant bioinformatics workflows.

In addition to data-layer control, OpsAgent adopts a namespaced project model that enables secure operations and collaborations. Projects function as isolated workspaces where users, tasks, and data objects are logically grouped, with OpsAgent coordinating membership, task assignment, and data bindings. This coordination layer enforces consistent interface contracts and guarantees that downstream analytical agents operate on schema-conformant, semantically validated inputs. It also supports fine-grained permission settings through role-based access control (RBAC), allowing users to be assigned scoped privileges—such as read, write, or administrative access—ensuring that collaborative activities occur within clearly defined security boundaries. By coupling strict input governance with robust access control, OpsAgent facilitates secure, reproducible, and scalable execution across multi-agent pipelines while supporting seamless collaboration among users.

### 2.3 DataAgent

DataAgent streamlines data discovery and acquisition for bioinformatics analyses by addressing two key user needs:

1. Effective retrieval of relevant data from public datasets through natural language query.
2. Seamless access to retrieved data via efficient cloud-to-cloud transfer.

The goal is to comprehensively and accurately identify the most relevant publicly available datasets based on user queries, while also assessing whether the data is directly downloadable. For datasets with controlled access, the system allows users to provide access credentials or tokens, enabling secure and straightforward data import.

DataAgent integrates a broad range of specialized repositories that host, manage, and annotate diverse types of biological and genomic data. These curated sources are critical to advancing scientific research by providing structured and reliable access to large-scale datasets. A full list of supported databases is provided in Supplementary Table 1, covering the majority of commonly used omics data types and repositories. This list is readily extensible to incorporate additional data sources as needed.

#### 2.3.1 Query Processing and Semantic Search

DataAgent streamlines the dataset search process by combining natural language understanding with structured search capabilities. Large Language Models (LLMs) parse user queries (e.g., “*I would like to obtain RNA-seq data for Alzheimer’s disease with a sample size greater than 30*”) and extract both the core search intent (e.g., “*RNA-seq data for Alzheimer’s disease*”) and specific filtering criteria (e.g., “*sample size greater than 30*”). A hybrid search engine then integrates semantic similarity—matching user intent to dataset descriptions in metadata—with keyword-based retrieval and metadata-aware filtering to efficiently surface the most relevant datasets.

To assess DataAgent’s retrieval accuracy, we conducted a benchmark study, detailed in the **Supplementary Table 2**. The results demonstrate its effectiveness in returning datasets that align closely with both the semantic content and specific conditions of user queries.

#### 2.3.2 Efficient Cloud-to-Cloud Data Transfer

DataAgent enables direct cloud-to-platform data transfers from AWS S3 and other cloud-hosted data repositories, bypassing the burden of transferring large files through local machines. By automating authentication and leveraging secure repository APIs, DataAgent ensures both speed and data security. It is also equipped with fault-tolerant features such as automatic transfer resumption to handle transient network interruptions. All retrieved data and metadata are ingested into the PromptBio data system, making them immediately available for downstream analysis. Designed for scalability, DataAgent supports diverse omics data types—enabling streamlined, secure integration across a broad range of research applications.

### 2.4 OmicsAgent

OmicsAgent delivers standardized, scalable solutions for omics data processing and analytical workflows. The platform supports a comprehensive suite of pipelines for multi-omics data types and commonly used downstream analysis modules. Each solution is optimized for performance and adherence to domain-specific best practices, ensuring reproducibility. For a detailed technical overview of the supported omics pipelines, analytical modules, and implementation details, refer to Section 3.1 and 3.2.

### 2.5 AnalysisAgent

AnalysisAgent is designed to meet flexible, custom analytical needs for tabular data that fall outside the scope of standard OmicsAgent functionality. It empowers researchers with intelligent, on-demand analytics through two complementary modes of interaction, each optimized for different levels of complexity and transparency.

- **Fast Analysis Mode** is tailored for rapid prototyping and lightweight, results-oriented tasks. Users can submit queries in natural language—such as “*What is the dimension of my input data matrix?*” or “*Create a scatter plot of the gene expression values of EGFR vs. PIK3CA*”—and receive responses from automatically generated Python or R scripts execution results. The outputs include summary, visualizations, and concise interpretations that provide immediate insight without requiring manual coding.
- **Notebook Mode** supports more complex, exploratory workflows where transparency and reproducibility are critical. In response to detailed prompts, AnalysisAgent generates well-structured Jupyter notebooks with clean, annotated code, embedded results, and interpretations. For example, a user might request: “*Merge data1.csv and data2.csv, apply batch effect correction, compare the PCA results before and after correction.*” These notebooks are fully executable within the PromptBio environment, enabling users to inspect, modify, rerun, and share analytical workflows with ease.

By uniting intelligent code synthesis, contextual understanding, and automatic execution, AnalysisAgent ensures that researchers can efficiently obtain actionable insights tailored to their specific scientific questions.

### 2.5 QAgent

QAgent is designed to streamline user interactions and enhance the bioinformatics analysis experience. By combining Retrieval-Augmented Generation (RAG) with large language models (LLMs), QAgent delivers accurate, context-aware responses while maintaining strict data privacy where each user is restricted for queries under his own workspace.

QAgent addresses three main types of questions using distinct information sources (Figure 2):

- **General knowledge questions** are answered using the LLM’s own knowledge base (e.g., “*What is the TP53 gene?*”).
- **Platform-related questions** are resolved by referencing curated documentation from a vector database, providing detailed guidance on functionalities, tool options, and usage instructions (e.g., “*What is the PromptBio genomics pipeline?*”). An example is shown in **Supplementary Figure 1**.
- **User workspace questions** are addressed by accessing relevant workspace information, such as project or task identifiers, to extract findings or results (e.g., “*What are the key findings of the differential analysis for task ID …?*”). An example is shown in **Supplementary Figure 2**.

**Figure 2:**
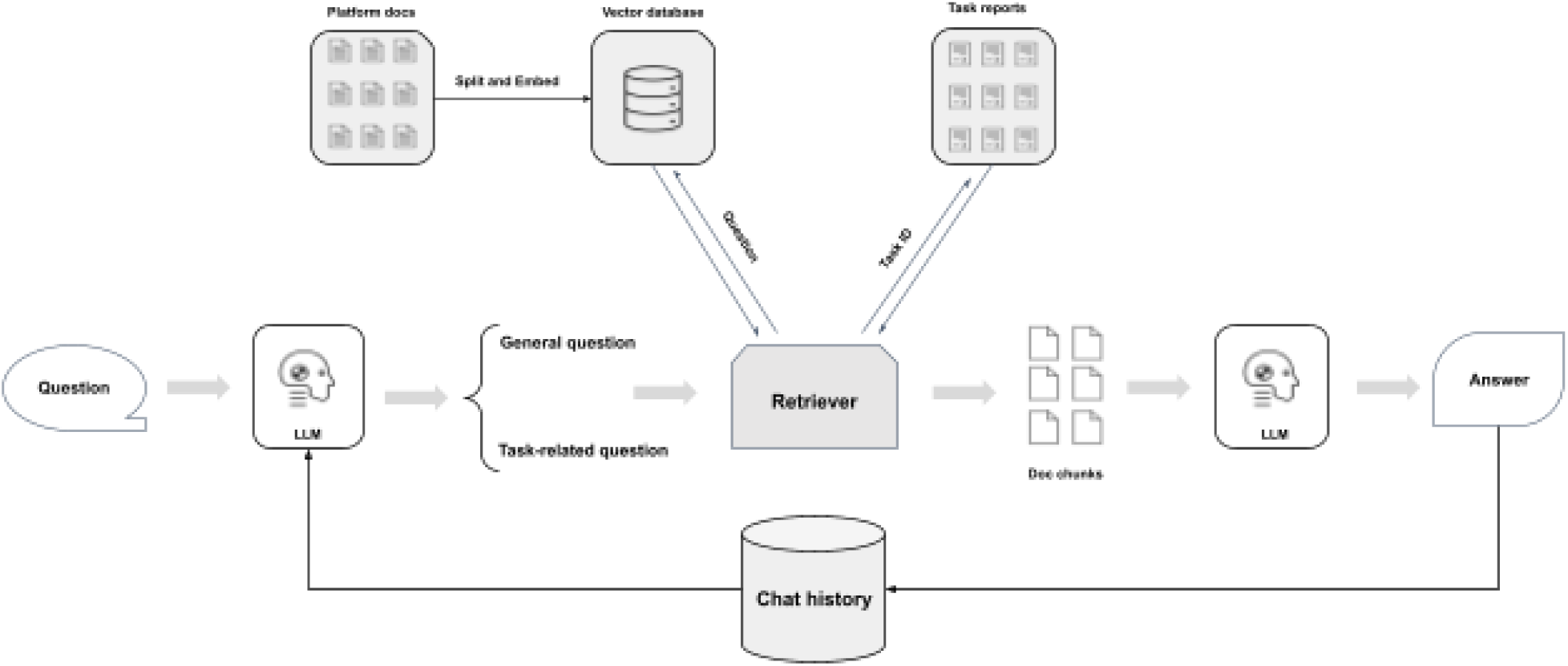
The design of QAgent.

The selected information is processed by the LLM to generate precise, conversational responses, with chat history continuously updated to ensure context-aware interactions. By integrating these three information sources, QAgent enables natural language interaction with PromptBio—covering platform features, tool settings, task status, and analysis results. This approach reduces reliance on manuals and support tickets, making the platform more accessible and user-friendly for both novices and experts, while providing clear explanations and result interpretations to support informed decision-making.

## 3 Expansive Suite of Tools

The PromptBio platform offers a comprehensive suite of tools, supporting both routine and advanced bioinformatics analysis for omics data. The suite integrates community-standard **omics tools and pipelines** (e.g., Seurat, GATK) with proprietary modules such as **MLGenie** for statistical modeling and **MarkerGenie** for biomarker discovery. This tool suite enables extensible omics analyses, reducing manual effort while supporting a wide range of research needs.

### 3.1 Omics Tools

Omics Tools consist of standardized pipelines and analysis modules to process various omics data, such as genomics, transcriptomics, proteomics, and single-cell omics. Each module leverages well-established bioinformatics softwares and may involve multiple steps executed sequentially – from raw data quality control, preprocessing, to advanced statistical modeling and biological interpretation. These modules ensure analysis accuracy by adhering to best practices, significantly enhancing results quality and reproducibility.

#### 3.1.1 Omics Pipelines

Currently, the platform integrates several in-house, validated pipelines that support routine bioinformatics data processing across a broad spectrum of omics domains, including:

- **Genomics**: The genomics pipeline processes whole genome sequencing (WGS), whole exome sequencing (WES), and targeted sequencing data to comprehensively identify genomic alterations, including single nucleotide variants (SNVs), insertions and deletions (indels), copy number variations (CNVs), and structural variants (SVs). It further supports the analysis of key genomic biomarkers such as microsatellite instability (MSI) and tumor mutational burden (TMB), offering valuable insights into genetic variations, diagnostic biomarkers, and potential therapeutic targets.
- **Transcriptomics**: this pipeline processes RNA-seq data end-to-end: from raw sequencing data, encompassing gene expression quantification and fusion gene detection using state-of-the-art methods.
- **Proteomics**: this pipeline supports the analysis of mass spectrometry (MS)-based data, covering peptide identification, protein inference and quantification, and characterization of post-translational modifications (PTMs). The workflow is compatible with raw data formats from a broad range of mass spectrometry platforms—including Thermo Scientific, Bruker, Waters, Sciex, and Agilent—ensuring wide applicability.
- **Single-Cell Transcriptomics**: Tailored for high-resolution cellular level analysis, this pipeline supports data from platforms such as 10x Genomics, Parse Biosciences, and HIVE.
- **Methylomics**: this pipeline processes bisulfite sequencing data to deliver genome-wide DNA methylation profiling. Leveraging tools such as Bismark and BWA-Meth, the pipeline enables accurate detection and quantification of methylation patterns, supporting epigenetic studies in both research and clinical contexts.
- **Other Omics Applications**: Beyond these core areas, we offer specialized pipelines for emerging and niche omics applications, including T-cell receptor (TCR) profiling and metabolomics. Additional workflows support critical analyses such as human leukocyte antigen (HLA) typing, homologous recombination deficiency (HRD) assessment [13, 14], and neoantigen prediction, ensuring comprehensive multi-omics support tailored to diverse research and translational needs.

These in-house pipelines have been rigorously validated for accuracy and reproducibility through extensive internal benchmarking and comparison against established external tools (**Supplementary Data 5**), empowering researchers to uncover biologically meaningful insights with confidence.

#### 3.1.2 Analysis Modules

Complementing our primary processing pipelines, PromptBio integrates a suite of downstream analytical modules to support biological interpretation and insight generation from processed feature matrices—either generated by omics pipelines or directly uploaded by users, including:

- **Differential Analysis:** To streamline the identification of differentially expressed markers, PromptBio features a curated in-house module supporting diverse data types, including raw RNA-seq counts, binary mutation data, and normalized continuous data. Widely used methods such as DESeq2, EdgeR, Limma, Fisher’s exact test, Student’s t-test, and Wilcoxon test are implemented for rigorous statistical modeling and testing. Results can be visualized in various scientific figures including volcano, box, MDS, and MA plots, enabling intuitive interpretation and highlighting biologically meaningful changes.
- **Functional Enrichment Analysis:** PromptBio offers robust functional enrichment analysis to interpret differential marker changes and biological pathways. It supports both over-representation analysis and gene set enrichment analysis, utilizing curated databases such as Gene Ontology (GO), Kyoto Encyclopedia of Genes and Genomes (KEGG), and Reactome. Users can analyze gene lists or pre-ranked genes across various organisms and gene set categories, facilitating comprehensive understanding of underlying molecular mechanisms.
- **Multi-Group Comparison:** PromptBio supports both ordered and unordered group analyses. For ordered groups, methods such as Jonckheere’s trend test and Cochran–Armitage test are implemented. For unordered groups, ANOVA, Kruskal-Wallis, and Chi-squared tests are offered, accommodating both continuous and binary data. The analysis ensures data quality through normalization, missing value filtering, and feature selection. Multiple testing correction, including the Benjamini-Hochberg procedure, controls false discovery rates. Results are visualized with volcano plots, heatmaps, and MA plots for clear interpretation of molecular changes across groups.
- **Single-Cell Analysis:** PromptBio offers a comprehensive single-cell RNA-Seq clustering workflow, beginning with quality control, normalization, and identification of highly variable genes. It supports flexible clustering using methods such as Seurat, Monocle3, Scanpy, and RAPIDS (GPU-accelerated). Following clustering, marker genes for each group are identified using statistical tests like the Wilcoxon test, enabling detailed characterization of cellular subpopulations. Cell types are inferred from marker profiles using LLM-based methods for deeper biological insight. Results are compiled into detailed reports and visualized with dimensionality reduction techniques such as UMAP or t-SNE to reveal cellular heterogeneity.

These modules are fully compatible with upstream pipelines and have been validated for reproducibility and accuracy, enabling seamless, end-to-end omics analysis within the platform.

#### 3.1.3 Community-Standard Integration

To improve the accessibility of other advanced omics tools, we also integrated external peer-reviewed pipelines from the nf-core community [15], providing over 80 standardized workflows for applications such as *nf-core/genomeassembler* and *nf-core/ampliseq* to support additional research needs. For example, a recent multi-omics case study [16] integrated the in-house transcriptomics pipeline with *nf-core/ampliseq* to investigate host–microbiome interactions in cystic fibrosis, showcasing the platform’s versatility in supporting dynamic and diverse analysis needs for complex research questions.

However, while powerful, nf-core pipelines remain challenging for non-specialists due to difficulties in selecting appropriate pipelines, preparing tailored sample sheets, and configuring parameters. To overcome these barriers, we developed a recommendation module, which utilizes PromptGenie to interpret user queries and retrieve the most suitable nf-core pipeline based on official documentation and community discussions. The future release will enable automatically configuring tailored input metadata and reliable, context-aware parameter settings. Together, these features substantially minimize manual effort and lower the barrier for non-specialists, making nf-core pipelines more accessible.

### 3.2 MLGenie

MLGenie is a comprehensive machine learning module providing data preprocessing, feature selection, model training, and multi-omics integration. The module is organized into four core components—**Exploratory Data Analysis**, **Classification**, **Regression**, and **Survival Analysis**—that empower researchers to efficiently analyze any matrix data, such as high-dimensional omics data, and derive actionable insights.

Key features of MLGenie include:

- **Data Preprocessing:** MLGenie automates comprehensive data preprocessing to ensure high-quality, analysis-ready datasets. This includes imputation of missing values, normalization, and outlier detection. Specialized filters identify and retain only high-quality, informative features. Highly correlated features are removed to reduce redundancy and multicollinearity. Distinct workflows are applied to binary and continuous features, ensuring each is cleaned, imputed, and transformed appropriately.
- **Feature Selection:** MLGenie applies advanced techniques such as LASSO, mutual information, variance thresholding, ANOVA F-score, AUC, and correlation-based selection to reduce dimensionality and boost interpretability. Both univariate and model-based selectors, including recursive feature elimination, are supported for classification and regression. Statistical rigor is enhanced through bootstrap sampling, permutation testing, and FDR control. By integrating multiple approaches, MLGenie identifies the most informative features, improving both model accuracy and interpretability.
- **Model Selection and Optimization:** MLGenie streamlines model selection and optimization for classification, regression, and survival analysis. The system supports a wide range of algorithms—including random forests, gradient boosting, neural networks, support vector machines, and Cox models—tailored to specific tasks. Cross-validation and hyperparameter tuning algorithms are employed to identify the best-performing models, ensuring performance and model generalization. MLGenie allows customized model configuration, parallel processing and reproducible results, facilitating model comparison and optimization towards reliable machine learning outcomes.

The design of MLGenie ensures both scalability and adaptability, establishing it as a versatile, user-friendly tool for applications such as disease classification, biomarker discovery, and survival prediction.

### 3.3 MarkerGenie

MarkerGenie is a standalone tool originally developed to facilitate disease-specific biomedical entity relation extraction [17]. Its primary objective is to automate the identification and extraction of relationships—such as gene-disease associations—from unstructured biomedical literature, thereby providing a more efficient and less biased alternative to manual curation. We further improved the performance by integrating more advanced Named Entity Recognition (NER) techniques and the implementation of Shortest Dependency Path (SDP)-based negative training data filtering. The improved performance is provided in **supplementary data section 3**.

Leveraging MarkerGenie, users can systematically identify disease-associated biomarkers from public literature, with results ranked according to their frequency of appearance. The platform supports the discovery of a wide range of biomarkers, including genes, chemicals (such as metabolites), and microbiome entities. For example, in the context of endometrial cancer, MarkerGenie successfully identifies top-ranked biomarkers such as PTEN, TP53, 17α-Ethynylestradiol, and Atopobium vaginae—entities that are well-established in the literature as being associated with endometrial cancer (**Supplementary Table 4**).

MarkerGenie is now an integral component of the PromptGenie framework, and plays a pivotal role in facilitating the interpretation of results generated by other analytical modules, such as differential analysis and machine learning-based marker identification. By combining MarkerGenie’s literature-based search results with LLM-driven summarization, the system provides a comprehensive and context-rich interpretation of biomarker findings. In contrast to purely LLM-based approaches, MarkerGenie provides a more reproducible, transparent, and trackable methodology for biomarker validation, ensuring that supporting evidence can be obtained.

## 4 DiscoverFlow: A Framework for Automated Multi-Omics Data Analysis

PromptGenie enables step-by-step, human-in-the-loop analysis, but a key challenge remains: unifying diverse components into configurable and reproducible workflows. Ensuring consistent results across datasets requires streamlined workflow management and strong reproducibility guarantees.

To address this, we developed DiscoverFlow—a dedicated module for flexible, scalable, and automated end-to-end multi-omics analysis. We describe its system design, workflow creation, and workflow execution in the following sections.

### 4.1 Design

DiscoverFlow models data analysis workflows using a directed acyclic graph (DAG) framework, where each path in the graph represents a complete analytical process. In this structure, nodes correspond to either data inputs or processing steps, while edges represent the flow of data between them.

For example, in a typical multi-omics workflow (Figure 3): (1)Genomics, transcriptomics, and proteomics datasets serve as initial input nodes. (2) Each dataset is routed through its omics-specific processing pipeline. (3) The resulting processed feature matrices are then combined and passed to a classification node for downstream analysis.

**Figure 3:**
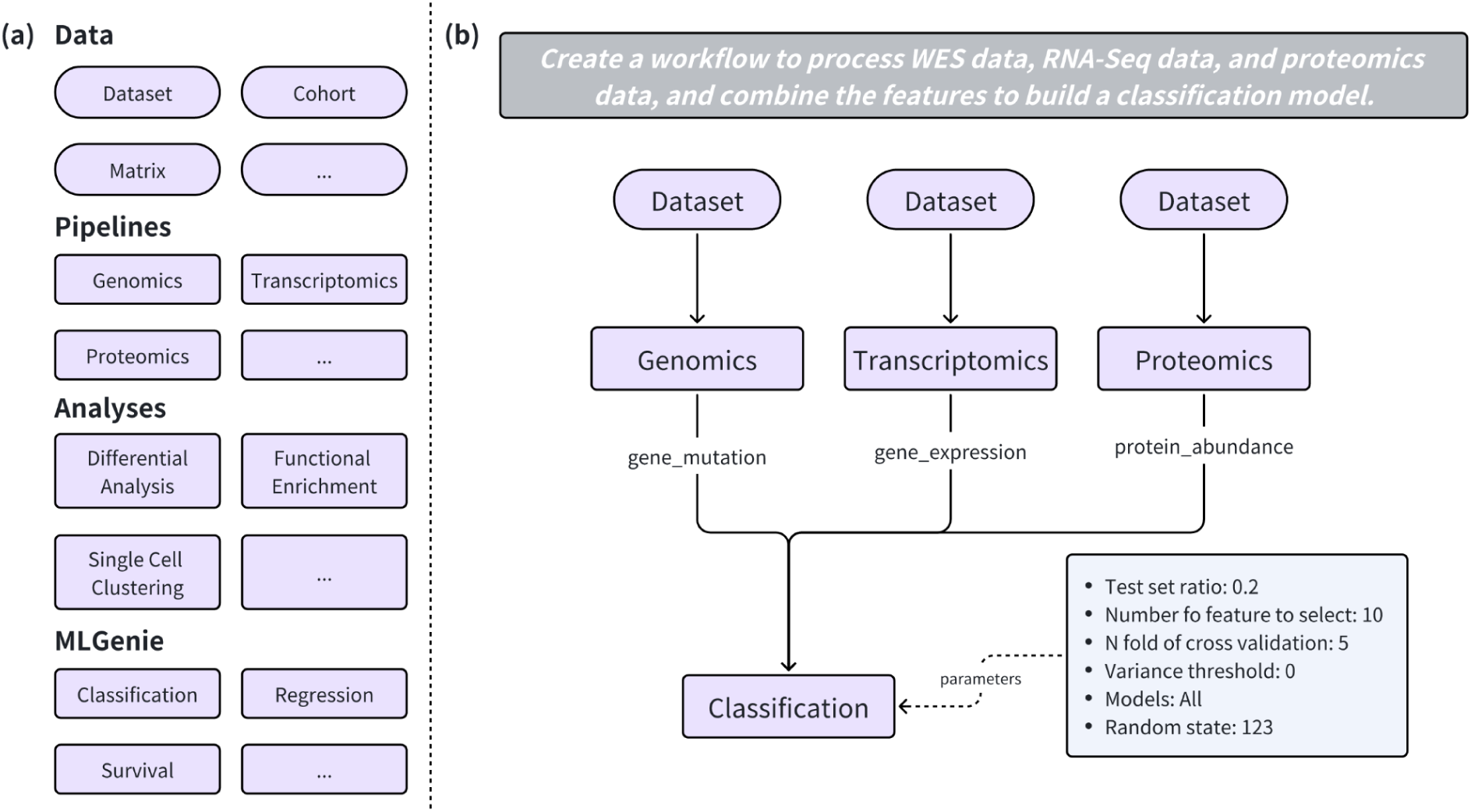
(a) Different building blocks currently available in DiscoverFlow. (b) Example workflow generated in response to the prompt: “Create a workflow to process WES data, RNA-Seq data, and proteomics data, and combine the features to build a classification model.”

The platform currently supports nodes spanning the full spectrum of omics workflows—including data preprocessing, downstream analysis, machine learning, and biomarker validation, as detailed in Section 3. Node dependencies are explicitly managed to enforce valid data flow—for example, a differential expression node can feed into a functional enrichment analysis, while incompatible connections (e.g., linking proteomics data directly to single-cell clustering) are disallowed. The node library is also designed for extensibility, enabling integration of new analytical modules.

### 4.2 Workflow Creation and Management

DiscoverFlow offers an intuitive, visual interface that interprets natural language input to help users construct, customize, and visualize tailored analysis workflows. Each workflow node can be customized, providing simplicity for beginners and fine-grained control for advanced users. This design lowers the barrier for those without deep bioinformatics expertise. Created workflows can be saved, shared, and re-executed—ensuring reproducibility and supporting collaborative research.

### 4.3 Workflow Execution

Task execution within DiscoverFlow is managed by a graph-based engine that automatically resolves dependencies and orchestrates parallel execution of independent tasks. Execution begins at the starting data node and progresses through the directed graph, with each node being processed as soon as all its dependencies are satisfied. It ensures efficient resource utilization and rapid turnaround for large-scale analyses.

In summary, DiscoverFlow unifies diverse analytical capabilities within a single, extensible platform, enabling the creation of configurable and reproducible workflows for end-to-end multi-omics data analysis. Its modular node-based design not only supports a comprehensive suite of current bioinformatics tasks but is also designed to accommodate future analytical innovations, ensuring ongoing adaptability and scalability for the evolving needs of the research community.

## 5 ToolsGenie: Enabling Custom, On-Demand Bioinformatics Analysis

ToolsGenie is developed to support custom workflows tailored to unique hypotheses, data modalities, or experimental designs that fall outside conventional pipelines. It is an AI-powered module built for dynamic, on-demand analysis.

Unlike fixed workflows in Omics Tools or DiscoverFlow, ToolsGenie enables users to construct entirely new pipelines from natural language prompts. The platform automatically selects appropriate methods, generates scripts, and assembles executable workflows, granting users complete freedom beyond the current tool suite provided by PromptBio. This empowers researchers to iterate rapidly, explore edge cases, and prototype novel analytic strategies.

### 5.1 Design

ToolsGenie is composed of four primary components (Figure 4), each fulfilling a distinct role in the workflow lifecycle:

- **Planner** The Planner acts as the system’s entry point, interpreting user analysis requests expressed in natural language and decomposes complex queries into logical, step-by-step analysis plans. This process ensures that each analytical objective is broken down into actionable subtasks, tailored to the user’s data type, research goals, and experimental design. Importantly, ToolsGenie allows users to review and provide feedback on the generated plan, enabling iterative refinement and ensuring the workflow aligns precisely with their scientific intent.
- **Generator** Building on the structured plan, the Generator selects appropriate bioinformatics tools and packages—such as GATK for whole-genome variant calling, Seurat for single-cell RNA-seq, or edgeR for differential expression—based on the requirements of each subtask. It then generates and customizes executable scripts in R, Python, or shell. When existing packages are insufficient, the Builder utilizes LLMs to synthesize novel code, ensuring flexibility for edge cases and novel analytic strategies.
- **Executor** The Executor is responsible for managing the computational environment and executing the assembled pipeline. It automates the installation and configuration of all required software and dependencies, primarily using Conda, to guarantee reproducibility and consistency. The Executor also handles parameter management, version tracking, and execution logging, enabling pipelines to be rerun on new datasets with minimal configuration.
- **Debugger** To support robust and iterative workflow development, the Debugger continuously monitors pipeline execution, captures error information, and facilitates troubleshooting. When errors occur, it can suggest or apply modifications to the pipeline, allowing users to resolve issues and optimize their analyses with little manual intervention.

**Figure 4:**
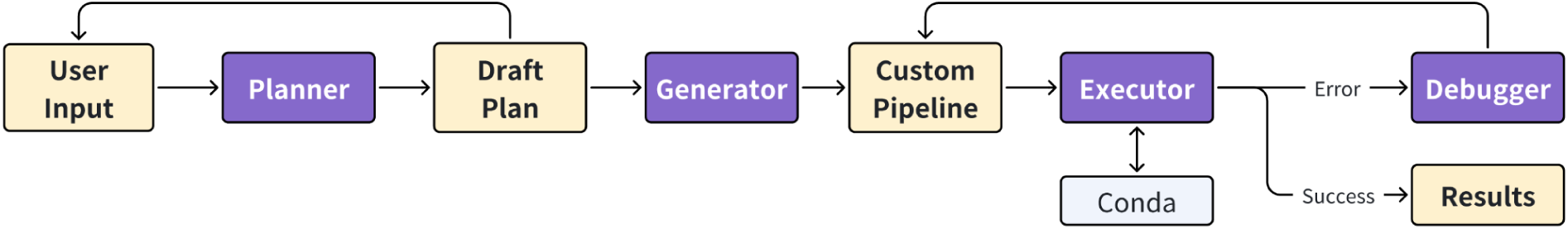
ToolsGenie Design. User input is first processed by the **Planner**, which generates a draft analysis plan. Upon user confirmation, the **Generator** refines this plan into a custom pipeline. The **Executor** then runs the pipeline, managing dependencies through Conda. If execution is successful, results are returned; otherwise, errors trigger the **Debugger**, which facilitates troubleshooting and pipeline modification prior to re-execution.

### 5.2 Application, Case Study and Benchmark

ToolsGenie is engineered to address the diverse and evolving analytical needs of modern biomedical research. Its flexible, AI-driven design enables users to design and execute workflows that range from simple data manipulations to complex, integrative analyses—without being constrained by pre-defined pipelines.

In a recent validation experiment, ToolsGenie was used to reproduce findings from the study “Interactions between the gut microbiome and host gene regulation in cystic fibrosis.” This research explores how host gene expression and gut microbiome composition interact in individuals with cystic fibrosis (CF).

With ToolsGenie, we processed RNA-seq data to identify genes that are differentially expressed in CF patients, revealing important pathways related to disease and host response. We also analyzed 16S rRNA sequencing data to profile the gut microbiome, uncovering changes in microbial diversity and abundance linked to CF. By integrating both data types, ToolsGenie helped uncover correlations between host gene activity and specific microbes, shedding light on the molecular interactions in the CF gut environment. Full details of this validation experiment are detailed in this [case study](https://d2v49qr4j6hcsp.cloudfront.net/PB-PDF/Multi-Omics%20Analysis%20of%20Cystic%20Fibrosis%20using%20Conversation-Based%20Interactions.pdf).

In addition to the case study, we conducted an internal benchmarking experiment involving 249 bioinformatics analysis tasks spanning a broad range of domains and difficulty levels. This evaluation assessed ToolsGenie’s ability to interpret natural language queries, select appropriate methods, and generate valid, executable workflows. The results—including accuracy metrics and detailed performance breakdowns—are presented in **Supplementary Section 5**.

### 5.3 Integration within the PromptBio Ecosystem

Although PromptGenie and ToolsGenie have been developed with distinct designs, ToolsGenie is natively integratable to the broader PromptBio multi-agentic framework. While there is currently some overlap in their functionalities, the ongoing refinement and expansion of the tool suite will enable ToolsGenie to play an increasingly pivotal role within the agentic ecosystem. This will allow for more seamless interaction with other agents and tools in the PromptGenie system. In this collaborative environment, ToolsGenie will be able to fully leverage the advanced data management capabilities of PromptBio, ensuring that workflows remain highly flexible and customizable, while also maintaining the robustness and reproducibility essential for scientific research.

## 6. Monitoring, Compliance, and Security

The PromptBio platform incorporates robust mechanisms for monitoring, regulatory compliance, and data security to ensure reliable and trustworthy bioinformatics analysis, particularly for sensitive patient data. These mechanisms are critical for clinical applications and large-scale deployments.

### 6.1 Monitoring

- **Real-Time Logging**: Captures workflow execution details, including task status, agent performance, and resource usage, centrally stored for streamlined access and troubleshooting.
- **Performance Metrics:** Continuously tracks critical performance indicators, including processing duration, memory consumption, and error rates, to facilitate proactive performance optimization.
- **Alerting:** Integrated notification systems promptly alert administrators to anomalies such as agent failures, enabling rapid intervention and system recovery.

### 6.2 Security

PromptBio employs advanced security protocols to protect data integrity and confidentiality:

- **Encryption:** Implements AES-256 encryption standards for data at rest and utilizes TLS protocols for secure data transmission, effectively safeguarding omics and other sensitive datasets.
- **Access Controls:** Incorporates role-based access control (RBAC) mechanisms, ensuring data access is strictly limited to authorized personnel only.
- **Secure APIs:** Secures agent and service communications through OAuth 2.0 authentication, enforcing strict control over system interactions and integrations.

### 6.3 Compliance Readiness

PromptBio is designed with a strong emphasis on regulatory compliance readiness, laying the groundwork for future adherence to major standards:

- **HIPAA** [22] **and SOC2** [23] **Preparedness:** Systems and processes have been structured to support patient data privacy through robust anonymization practices and secure storage strategies.
- **Audit Trails:** Maintains comprehensive records detailing data access, processing activities, and user actions, facilitating future compliance audits and regulatory assessments.

## 7 Discussion and Future Components

PromptBio demonstrates that large-scale, AI-driven bioinformatics analysis can be made more accessible, scalable, and reproducible through a modular, multi-agent system. At its core is a **supervisor–worker architecture**, where **PromptGenie**, the supervisor agent, coordinates a set of specialized agents — including DataAgent, OmicsAgent, and AnalysisAgent — to carry out structured tasks. While effective for guided execution, the system does not currently support full agentic behavior such as autonomous planning or peer-to-peer agent interaction.

Looking ahead, we aim to **incrementally evolve** PromptBio toward a more **agentic framework** [18], where agents can dynamically plan and adapt based on user intent and data context. However, this transition presents notable challenges. In particular, early trials show that autonomous agents often struggle to generate coherent, multi-step workflows [19] (**Supplementary data 6**). These limitations are especially pronounced in life sciences, where precision and reproducibility are essential. For now, a **human-in-the-loop** paradigm remains critical, and agent autonomy must be introduced cautiously, with strong constraints and validation.

To support this evolution, we are prioritizing several enhancements:

- **Unified Planning**: Merging **ToolsGenie** and **PromptGenie** into a single, integrated framework will streamline workflow generation, enabling seamless transition between user-defined and system-guided analyses.
- **Explainability Module**: Adding support for interpretable outputs (e.g., SHAP values for feature importance) will help users better understand model behavior and increase confidence in downstream decisions.
- **Interactive Visualization**: A real-time dashboard agent will allow users to explore results through interactive plots (e.g., PCA, t-SNE, UMAP), aiding both interpretation and exploratory data analysis.
- **Public Data Integration**: Incorporating large-scale processed datasets such as **GTEx** and **TCGA** will provide valuable reference points, improve context for user analyses, and accelerate discovery.
- **Expanded Omics Support**: Enhancing coverage of emerging data types—such as spatial transcriptomics, ATAC-seq, ChIP-seq, and Hi-C—will broaden the platform’s applicability to cutting-edge multi-omics research.
- **Pipeline Modularization**: We will continue expanding the library of analytical building blocks, enabling users and agents to compose flexible, reusable workflows tailored to specific scientific questions.

Future work will focus on integrating these enhancements and validating the platform at scale through extensive benchmarking on diverse, real-world datasets. This will be critical for ensuring robustness, generalizability, and readiness for broader research and translational applications.

## Acknowledgments

We thank our external trial users for their valuable feedback, which directly contributed to improving the platform.

## Supplementary Data sample

### 1. Public Database Repositories used in DataAgent

**Supplementary Table 1:** Database Summary in DataAgent. The databases incorporated 12 commonly used databases, with their corresponding number of datasets and samples shown in the table. For proteomics databases (PRIDE, iProX, SCP), the number of samples cannot be directly derived from file counts and thus is marked as NA.

| Database | Number of Datasets | Number of Samples |
| --- | --- | --- |
| SRA | 534,967 | 33,155,723 |
| GEO | 274,086 | 9,308,747 |
| ArrayExpress | 78,992 | 2,908,474 |
| PRIDE | 32,352 | NA |
| GDC | 5,475 | 1,102,448 |
| iProX | 4,382 | NA |
| MetaboLights | 1,826 | 338,638 |
| SCP | 832 | NA |
| TCGA | 514 | 222,964 |
| CPTAC | 187 | 23,091 |
| CPTAC-pancan | 185 | 19,282 |
| PDC | 176 | 11,941 |

### 2. Validation Results of DataAgent

To assess the relevance of DataAgent’s search results, we designed eight representative data acquisition queries, spanning across different omics domains with diverse biological contexts. For each query, the top 10 and top 25 returned datasets were manually reviewed and labelled by bioinformatics experts, the precision score (#truly-related datasets / #returned datasets) for each query are calculated, as shown in the following table. We acknowledge that precision is relatively low in some cases (e.g., Case 3 and Case 4), primarily due to retrieved datasets not fully meeting the specified criteria. This is an area with clear room for improvement.

**Supplementary Table 2:** Precision of DataAgent’s data retrieval process. The top 10 and top 25 search results are shown in eight test cases.

| Cases | Query | Top 10 | Top 25 |
| --- | --- | --- | --- |
| 1 | Find spatial transcriptome data of human lung adenocarcinoma | 0.6 | 0.32 |
| 2 | Find plasma proteomics related to Alzheimer's disease | 0.7 | 0.28 |
| 3 | Find single-cell RNA data related to PD-1/PD-L1 resistance human melanoma | 0.2 | 0.2 |
| 4 | Find mouse T-cell methylation data | 0.2 | 0.12 |
| 5 | Find ATAC-seq data in human acute myeloid leukemia patients (don't want cell line data) | 0.7 | 0.44 |
| 6 | Find human gut metagenomics of Parkinson's disease | 0.8 | 0.88 |
| 7 | Find human patient microRNA related to colorectal cancer metastasis | 0.6 | 0.56 |
| 8 | Find human gene expression data related to platinum response for cancer (sample size greater than 30) | 0.7 | 0.36 |
| Average |  | 0.563 | 0.395 |

### 3. MarkerGenie Performance Optimization

The following is a performance comparison between the original MarkerGenie [17] and its improved version. The updated MarkerGenie enhances biomedical entity identification by integrating advanced Named Entity Recognition (NER) models with traditional string-matching techniques. It also applies the Shortest Dependency Path (SDP) method to more effectively detect and remove incorrect negative training examples, improving relation extraction accuracy.

For a consistent evaluation, the raw training data were generated using the same method as in [17], and the same test set was used. Across all three relation types and evaluation metrics, models trained with the updated methodology consistently outperformed those trained on the original data.

**Supplementary Table 3:** AUC scores of binary relation classification using original and updated MarkerGenie. All datasets contain approximately a 60:40 ratio of positive to negative samples.

| Relation Type | Methodology | AUC(train) | AUC(validation) | AUC(test) |
| --- | --- | --- | --- | --- |
| Disease-Microbiome | Original | 0.835 | 0.844 | 0.761 |
|  | Updated | 0.951 | 0.948 | 0.879 |
| Disease-Gene | Original | 0.846 | 0.827 | 0.771 |
|  | Updated | 0.944 | 0.943 | 0.801 |
| Disease-Metabolite | Original | 0.8332 | 0.836 | 0.781 |
|  | Updated | 0.9372 | 0.947 | 0.832 |

**Supplementary Table 4:** Top 5 identified markers associated with endometrial cancer.

| No. | Genes | Chemicals | Microbiome |
| --- | --- | --- | --- |
| 1 | PTEN | 17α-Ethinylestradiol | Atopobium vaginae |
| 2 | TP53 | Tamoxifen | Porphyromonas sp. |
| 3 | PIEZO1 | Progesterone | Porphyromonas |
| 4 | INS | 1,1-Dimethylbiguanide | Atopobium |
| 5 | MLH1 | Cisplatin | Porphyromonas somerae |

### 4. QAgent Examples

**Supplementary Figure 1:**
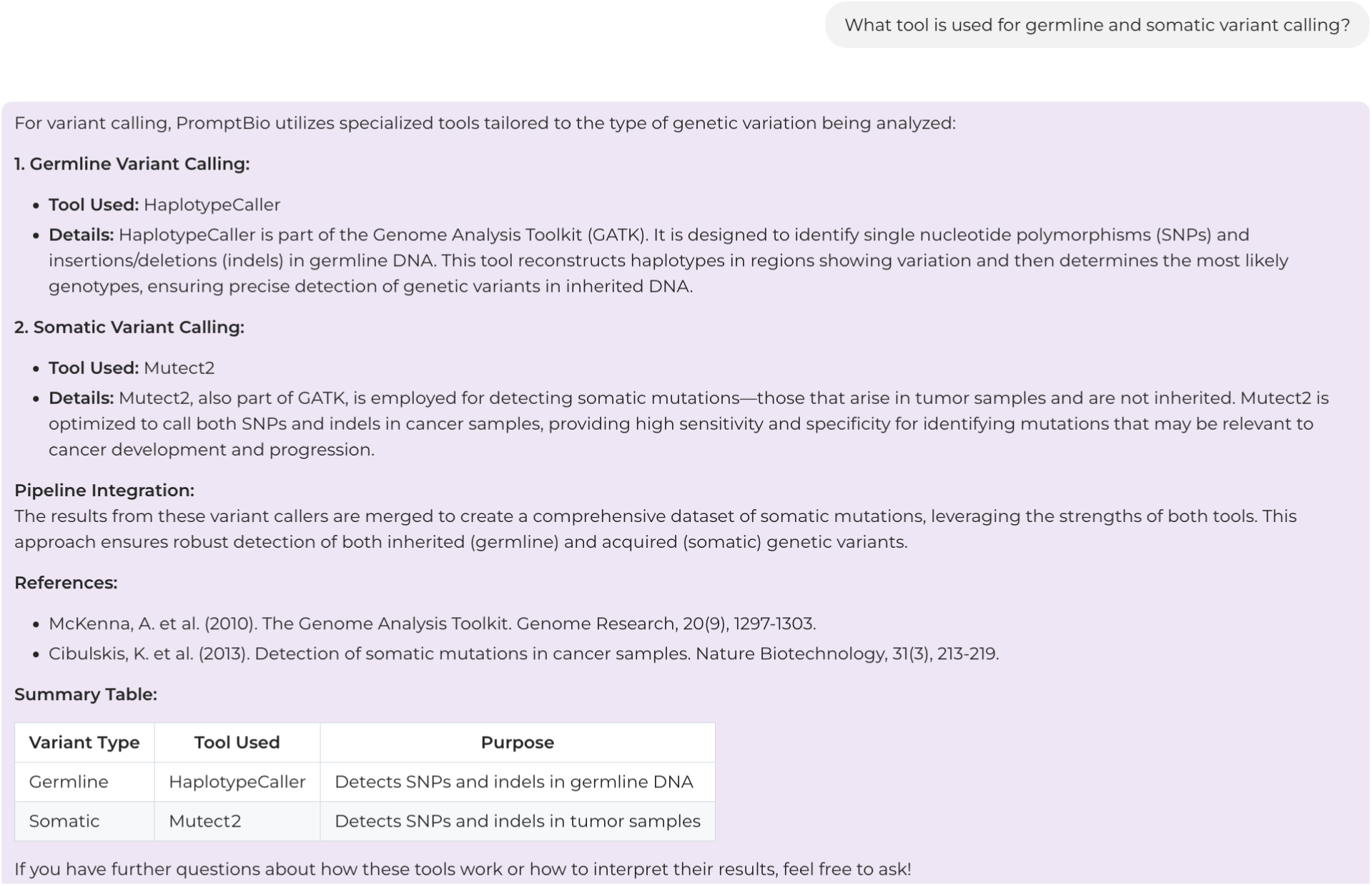
Example of system-related QA interaction.

**Supplementary Figure 2:**
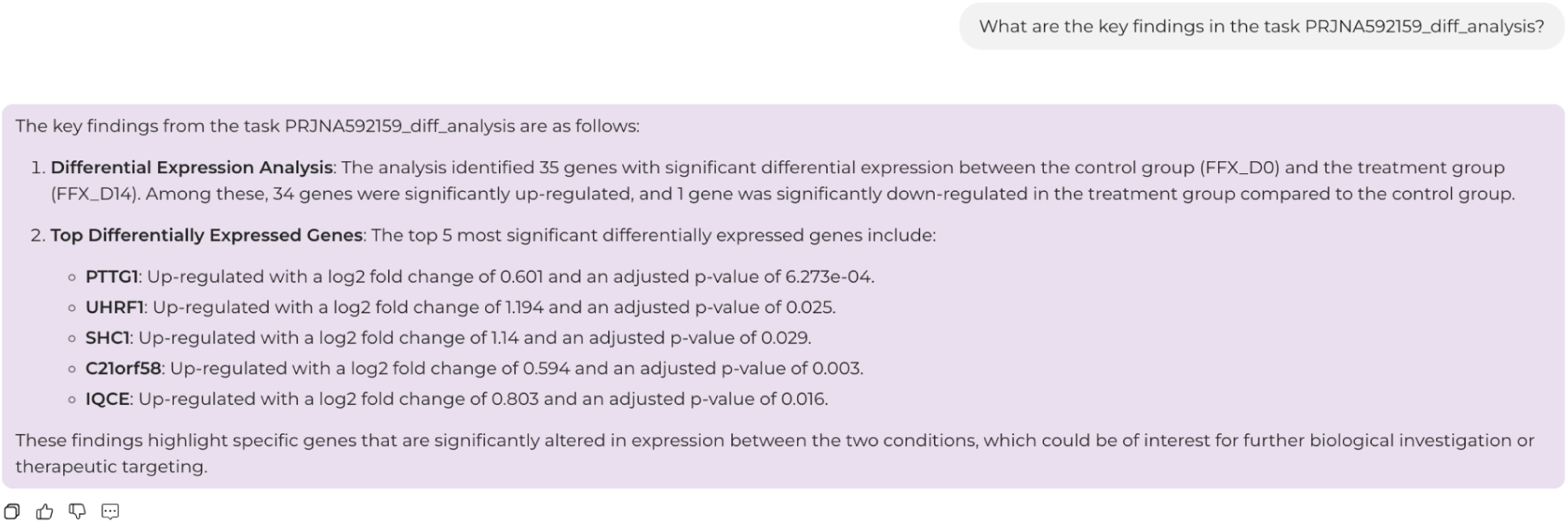
Example of result interpretation.

### 5. Omics Tools Validation Results

#### 5.1 Validation of Genomics Pipeline

We benchmarked both germline and somatic variant calling pipelines using external datasets. Specifically, we randomly choose a sample from Genome in a Bottle (GIAB) consortium and Sequencing Quality Control Phase 2 (SEQC2) consortium for germline variant calling and somatic variant calling, respectively. The results **(Supplementary Table 5**) show the in-house genomics pipeline (both CPU and GPU version) achieved comparable precision and recall than what the widely used Sarek pipeline (v3.5.1) did on both germline SNP calling and INDEL detection. While it’s suboptimal for somatic mutations, the difference is minimal, and the pipeline still demonstrated consistent and reliable performance.

**Supplementary Table 5:** Validation of germline and somatics variant calling of the Genomics Pipeline.

| Mutation type | Pipeline name | SNP |  |  | INDEL |  |  |
| --- | --- | --- | --- | --- | --- | --- | --- |
|  |  | Precision (↑) | Recall (↑) | F1 score (↑) | Precision (↑) | Recall (↑) | F1 score (↑) |
| germline | Sarek (v3.5.1) | 0.9662 | 0.9862 | 0.9761 | 0.9216 | 0.9133 | 0.9174 |
|  | PromptBio (CPU) | 0.9672 | 0.9862 | 0.9766 | 0.9222 | 0.9133 | 0.9177 |
|  | PromptBio (GPU) | **0.9845** | **0.9873** | **0.9859** | **0.9545** | **0.9164** | **0.9351** |
| somatic | Sarek (v3.5.1) | 0.9890 | 0.8568 | 0.9182 | **0.7167** | 0.8958 | 0.7963 |
|  | PromptBio (CPU) | 0.9941 | **0.8663** | **0.9258** | 0.6769 | 0.9167 | 0.7788 |
|  | PromptBio (GPU) | **0.9979** | 0.8361 | 0.9099 | 0.7031 | **0.9375** | **0.8036** |

#### 5.2 Validation of Proteomics Pipeline

Both protein identification and quantification were benchmarked using well-established benchmark datasets. Specifically, we used a yeast–UPS1 standard dataset acquired via LC-MS/MS (PRIDE database accession: PXD001819) to validate protein identification. We compared the set of proteins identified by our pipeline against those leading software tools, including MaxQuant (Intensity mode and LFQ mode), Skyline, and MFPaQ. The results showed a high degree of overlap, confirming the consistency and accuracy of our identification process (**Supplementary Figure 3**).

For protein quantification, we evaluated our pipeline using a real-world clinical dataset [20]. Specifically, we conducted a case study based on the proteogenomic dataset from the Cell article “Proteogenomic analysis of chemo-refractory high-grade serous ovarian cancer” (accessible via the PDC database). The protein expression results produced by our pipeline were compared against the article-reported expression profiles using Pearson correlation coefficients. As shown in **Supplementary Table 6**, over 90% of the samples in both frozen and FFPE validation cohorts achieved correlation coefficients greater than 0.8, indicating strong consistency between our results and those from the publication.

A detailed correlation matrix is presented in **Supplementary Figure 4**, where the FFPE validation cohort that consists of 20 FFPE samples are compared between PromptBio and the PDC reference data.

**Supplementary Figure 3:**
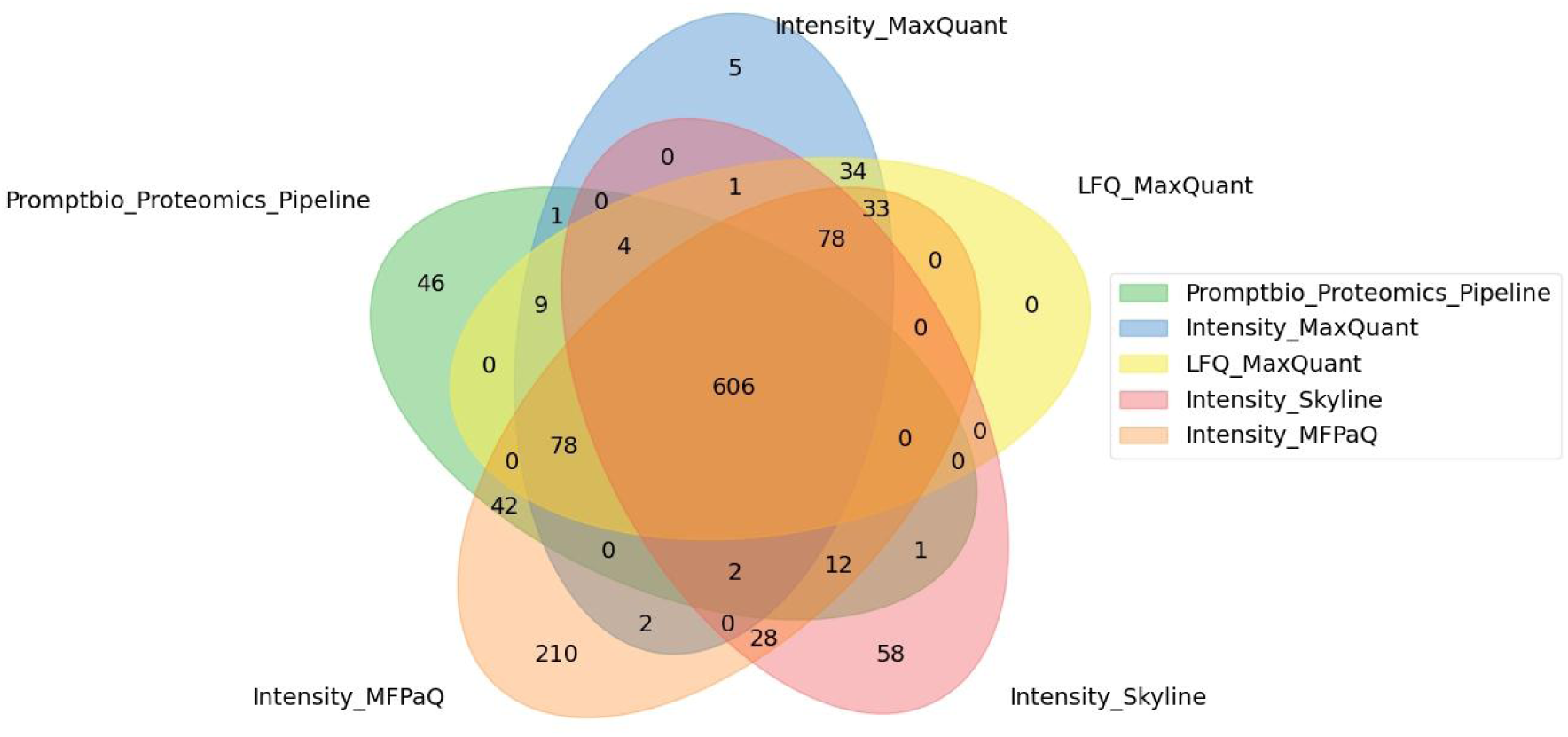
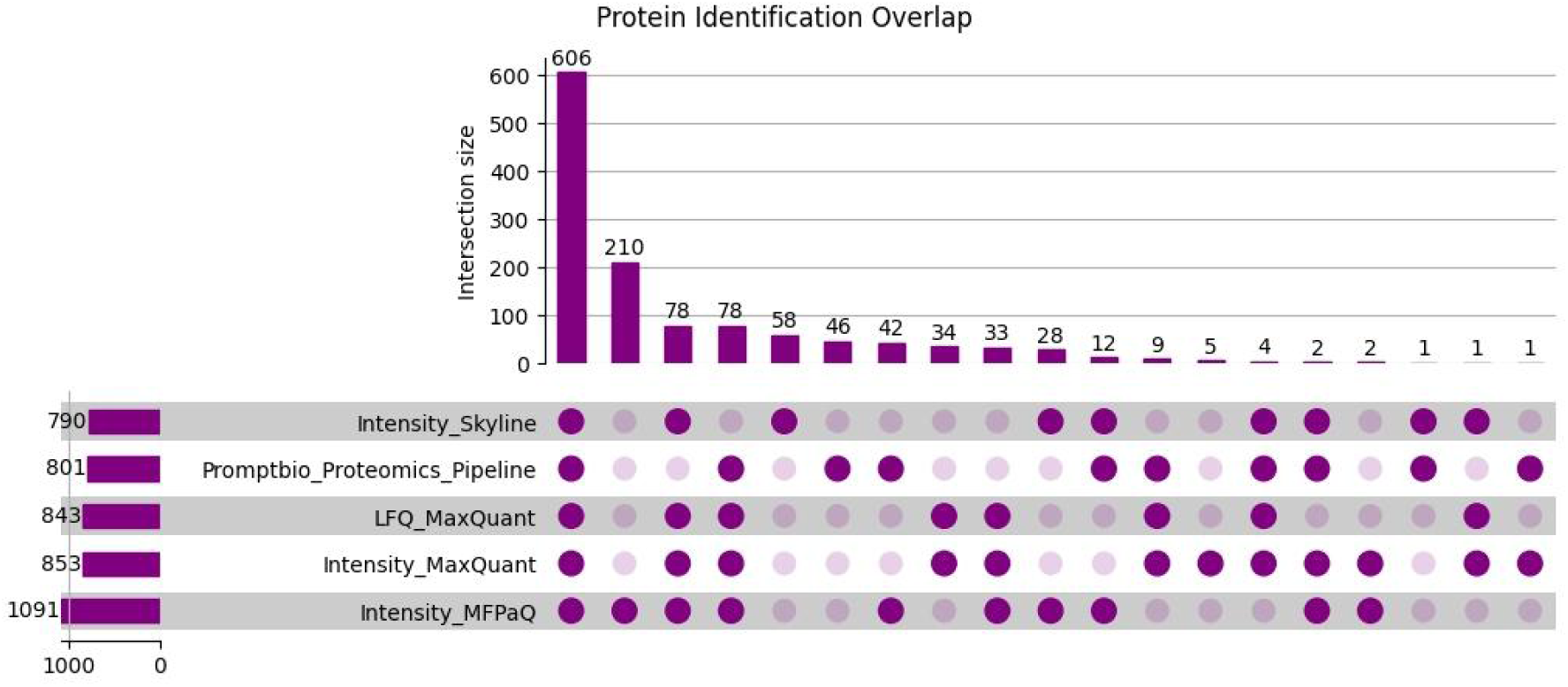
Validation of Protein Identification and Quantification. (Upper panel) Venn diagram showing the overlap of identified proteins between our pipeline and three external tools: MaxQuant, Skyline, and MFPaQ. (Lower panel) UpSet plot presenting the same overlap data as in the upper panel, providing a detailed view of shared and unique protein sets.

**Supplementary Table 6.** Correlation of protein expression levels between PromptBio and the Cell article. The proportions of samples where the correlation coefficient (R) is greater than a certain value are shown.

| Cohort | $R \geq 0.9$ | $R \geq 0.8$ | $R \geq 0.7$ |
| --- | --- | --- | --- |
| FFPE_Discovery_Cohort | 44.3% | 93.3% | 97.6% |
| Frozen_Validation_Cohort | 51.4% | 100.0% | 100.0% |
| FFPE_Validation_Cohort | 75.0% | 100.0% | 100.0% |

**Supplementary Figure 4.**
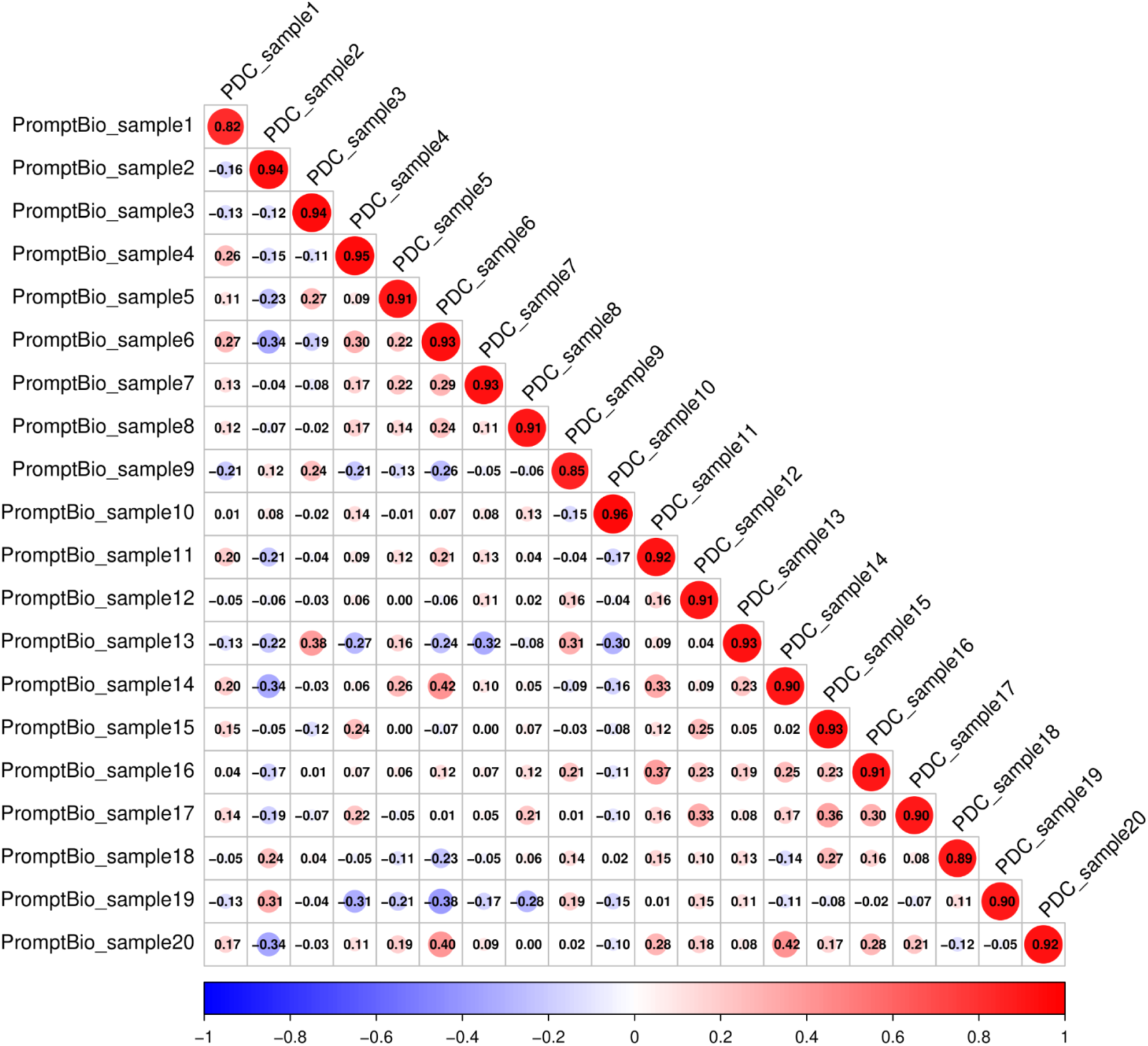
Correlation matrix plot comparing protein expression levels between PromptBio results and PDC data for the FFPE validation cohort (n = 20). Each cell represents a Pearson correlation coefficient between a sample pair, colored by correlation level.

### 6. ToolsGenie Validation Results

To evaluate the performance of **ToolsGenie** in addressing typical bioinformatics analysis tasks, we designed and conducted a targeted benchmarking study. A total of 259 representative questions were assembled, covering a broad range of bioinformatics domains and varying levels of analytical complexity. The questions were drawn from three main sources: (i) an in-house curated set of manually written prompts; (ii) newly generated questions based on data from the publicly available **BixBench** benchmark [21]; and (iii) structured questions derived from vignettes of widely used **CRAN R** packages, where representative code examples were translated into natural language prompts. This diverse question set was designed to reflect realistic use cases. Examples are presented in **Supplementary Table 7**.

For each question, we manually prepared the corresponding input data and constructed reference scripts, which were executed to generate ground truth answers. ToolsGenie was then tasked with solving each question using the same input data. Its outputs were systematically compared to the reference results. This evaluation was conducted manually, and an accuracy score was assigned to each response based on its alignment with the ground truth.

The assessment yielded an overall average accuracy score of **0.46** across all 259 questions. A detailed performance breakdown by sub-domain is presented in **Supplementary Table 8**, and common failure modes are summarized in **Supplementary Table 9**. Further analysis revealed that ToolsGenie performed reliably on straightforward, low-level tasks, particularly when the problem statement was clear and required minimal dependency resolution. In contrast, performance declined on questions involving multi-step analytical workflows, deeper contextual reasoning, or less common tools and packages. This is in parallel consistent with a separate benchmark effort applied to CRM domain [19], highlighting the significant challenges and opportunities.

These results underscore both the current strengths and limitations of ToolsGenie and provide important direction for future refinement and optimization.

**Supplementary Table 7.** Example benchmark questions by category and difficulty in ToolsGenie evaluation.

| Query | Category | Difficulty |
| --- | --- | --- |
| Filter the bam file with quality score threshold of 30 | NGS Processing | 1 |
| Perform hierarchical clustering on the dataset, split samples into two clusters, and visualize the dendrogram with cluster labels (colored by group) | Machine Learning | 2 |
| Perform a co-expression network analysis on the provided gene expression data and identify the key hub genes. | Bioinformatics Analysis | 3 |

**Supplementary Table 8.**
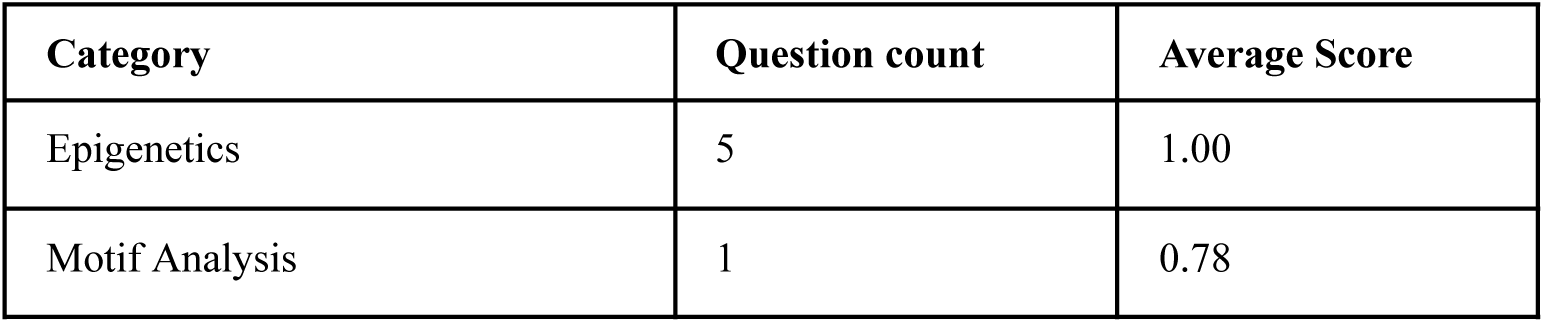
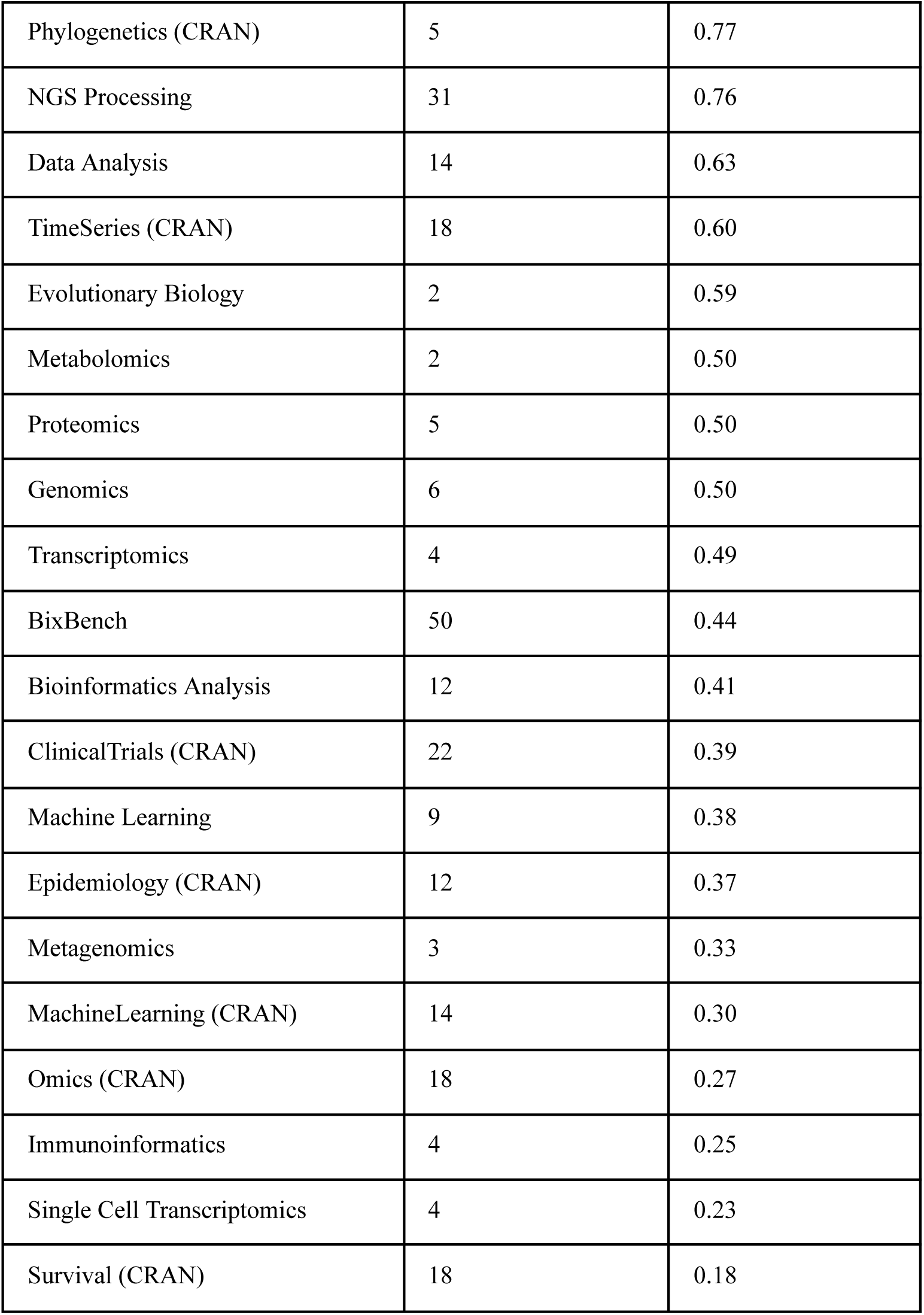
Average accuracy scores by analysis category in ToolsGenie benchmark evaluation. Categories labeled with “CRAN” indicate that the questions were derived from R CRAN resources.

| Category | Question count | Average Score |
| --- | --- | --- |
| Epigenetics | 5 | 1.00 |
| Motif Analysis | 1 | 0.78 |
| Phylogenetics (CRAN) | 5 | 0.77 |
| NGS Processing | 31 | 0.76 |
| Data Analysis | 14 | 0.63 |
| TimeSeries (CRAN) | 18 | 0.60 |
| Evolutionary Biology | 2 | 0.59 |
| Metabolomics | 2 | 0.50 |
| Proteomics | 5 | 0.50 |
| Genomics | 6 | 0.50 |
| Transcriptomics | 4 | 0.49 |
| BixBench | 50 | 0.44 |
| Bioinformatics Analysis | 12 | 0.41 |
| ClinicalTrials (CRAN) | 22 | 0.39 |
| Machine Learning | 9 | 0.38 |
| Epidemiology (CRAN) | 12 | 0.37 |
| Metagenomics | 3 | 0.33 |
| MachineLearning (CRAN) | 14 | 0.30 |
| Omics (CRAN) | 18 | 0.27 |
| Immunoinformatics | 4 | 0.25 |
| Single Cell Transcriptomics | 4 | 0.23 |
| Survival (CRAN) | 18 | 0.18 |

**Supplementary Table 9.** Common error types and their frequencies in ToolsGenie benchmark evaluation.

| Error Type | Count | Reason |
| --- | --- | --- |
| Misunderstanding Error | 42 | ToolsGenie misinterpreted the question's intent, lacked the necessary domain knowledge, or considered the input files insufficient. Causes include ambiguous or lengthy queries, or simple miscomprehension. |
| File Content Error | 19 | ToolsGenie was unable to correctly interpret input file contents, often due to unsupported file types (e.g., FASTA, RDS) or data matrices with excessive columns. |
| Misuse Package Error | 15 | ToolsGenie incorrectly used software packages, such as calling inappropriate functions. |
| Package Install Error | 12 | ToolsGenie failed to install required software packages. |

